# Display of malaria transmission-blocking antigens on chimeric duck hepatitis B virus-derived virus-like particles produced in *Hansenula polymorpha*

**DOI:** 10.1101/595538

**Authors:** David Wetzel, Jo-Anne Chan, Manfred Suckow, Andreas Barbian, Michael Weniger, Volker Jenzelewski, Linda Reiling, Jack S Richards, David A Anderson, Betty Kouskousis, Catherine Palmer, Eric Hanssen, Gerhard Schembecker, Juliane Merz, James G Beeson, Michael Piontek

## Abstract

1.

**Background:** Malaria caused by *Plasmodium falciparum* is one of the major threats to human health globally. Despite huge efforts in malaria control and eradication, highly effective vaccines are urgently needed, including vaccines that can block malaria transmission. Chimeric virus-like particles (VLP) have emerged as a promising strategy to develop new malaria vaccine candidates.

**Methods:** We developed yeast cell lines and processes for the expression of malaria transmission-blocking vaccine candidates Pfs25 and Pfs230 as VLP and VLP were analyzed for purity, size, protein incorporation rate and expression of malaria antigens.

**Results:** In this study, a novel platform for the display of *Plasmodium falciparum* antigens on chimeric VLP is presented. Leading transmission-blocking vaccine candidates Pfs25 and Pfs230 were genetically fused to the small surface protein (dS) of the duck hepatitis B virus (DHBV). The resulting fusion proteins were co-expressed in recombinant *Hansenula polymorpha* (syn. *Pichia angusta, Ogataea polymorpha*) strains along with the wild-type dS as the VLP scaffold protein. Through this strategy, chimeric VLP containing Pfs25 or the Pfs230-derived fragments Pfs230c or Pfs230D1M were purified. Up to 100 mg chimeric VLP were isolated from 100 g dry cell weight with a maximum protein purity of 90 % on the protein level. Expression of the Pfs230D1M construct was more efficient than Pfs230c and enabled VLP with higher purity. VLP showed reactivity with transmission-blocking antibodies and supported the surface display of the malaria antigens on the native VLP.

**Conclusion:** The incorporation of leading *Plasmodium falciparum* transmission-blocking antigens into the dS-based VLP scaffold is a promising novel strategy for their display on nano-scaled particles. Competitive processes for efficient production and purification were established in this study.

## 2. Background

Malaria is one of the world’s deadliest human diseases with nearly half of the global population living at risk. There were an estimated 216 million cases and 445,000 deaths due to malaria in 2016 [1]. This life-threatening disease is caused by *Plasmodium* parasites and is transmitted via the bite of infected female *Anopheles* mosquitoes. The majority of malaria is caused by *P. falciparum*, with *P. vivax* being a second major cause of disease [1]. Despite substantial financial investment, US$ 2.7 billion in 2016, and decades of intense research and development, only one malaria vaccine has progressed through phase 3 clinical trials and is now undergoing phase 4 implementation trials (RTS,S; Mosquirix^TM^). However, vaccine efficacy in phase III clinical trials was low in young children (up to 50% efficacy in the first year, but waning over 18 months) [2]. The World Health Organization has set a strategic goal of developing vaccines with at least 75% efficacy [3], including the development of vaccines that block malaria transmission [1]. Various approaches are under investigation including whole parasite vaccines and subunit vaccines that are composed of defined, purified antigens or their sub-domains [4]. Subunit vaccines have the potential to use established technologies and processes for low-cost production and distribution through existing vaccine delivery mechanisms [5]. A variety of *Plasmodium* antigens are currently under investigation as potential subunit vaccine components and can be classified into one of the following groups based on *Plasmodium* lifecycle stages [6]: i) pre-erythrocytic antigens (e.g. CSP [7]); ii) blood-stage antigens [8]; iii) transmission-stage antigens (e.g. Pfs25, Pfs230 [9–11]).

Unfortunately, subunit vaccine candidates often suffer from weak immunogenicity that has to be compensated by smart formulation and/or delivery strategies [12] such as virus-like particles (VLP [13, 14]). Since the 1980’s, VLP have been approved for use as safe and effective subunit vaccines against several pathogens [15]. They can also be used as a scaffold for the incorporation of antigens derived from foreign pathogens to enhance their immunogenic potential (chimeric VLP [16]). Accordingly, the RTS,S vaccine contains chimeric VLP with a truncated construct of CSP, the major surface antigen expressed on sporozoites during the pre-erythrocytic stage. However, its efficacy was low in young children; approaches are urgently needed to develop highly efficacious vaccines to improve malaria control and elimination [2, 17], such as the inclusion of additional antigens.

Recently, there has been a renewed focus of malaria vaccine development on transmission stage antigens, and transmission-blocking activity is a stated priority in the WHO Malaria Vaccine Technology Roadmap [3]. Transmission-blocking vaccines (TBV) are thought to act by inhibiting the transmission of malaria from humans to the mosquito vector, largely through the action of antibodies taken up in the mosquito’s blood-meal [18]. Leading vaccine candidates that are expressed during the transmission stages of *P. falciparum* include Pfs25 [9] and Pfs230 [10, 19]. Both antigens have been shown to generate antibodies that are capable of blocking transmission through standard membrane feeding assays [20–22]. Pfs230 is expressed on gametocytes and gametes and is a target of naturally-acquired antibodies from malaria-exposed populations, whereas Pfs25 is only expressed by zytgotes and ookinetes in the mosquito stage and, therefore, naturally-acquired immunity is not generated [18]. However, the development of TBV remains challenging.

The recombinant production and folding of Pfs25 is difficult because it is cysteine-rich and contains four tandem epidermal growth factor (EGF)-like domains [21]. Nevertheless, success has been achieved with yeast-derived versions of Pfs25 that are emerging as prominent TBV candidates [23–25]. However, the immunogenicity of Pfs25 is weak [26], but can be enhanced by construction of fusion proteins [27–29] or by VLP or non-VLP nanoparticle-based approaches [30–35]. The 363 kDa Pfs230 is a large and complex protein that is predicted to contain multiple cysteine-rich domains [36, 37]. Its potential as a transmission-blocking vaccine candidate was identified in the 1980s [10, 19]. However, recombinant production of full-length Pfs230 has not been accomplished to the present time. Therefore, research has focused on truncated Pfs230 versions named Pfs230c [22] and Pfs230D1M [38]. These variants were shown to retain the property to elicit transmission-blocking antibodies and can be produced as recombinant antigens [22,38,39]. The Pfs230c construct contains the first two protein domains. Expression studies of Pfs230 in yeast led to the development of the shortened Pfs230D1M construct, which includes only the first domain, and could be efficiently expressed in *Pichia pastoris* [38].

Despite progress that has been made towards effective malaria vaccines, novel protein conjugation strategies and delivery platforms as well as the use of strong adjuvants may be essential to meet the goals of the malaria control and eradication agenda set by the World Health Organization [40]. Our present study introduces a new platform for the display of the malaria transmission-blocking vaccine candidates Pfs25 [25], Pfs230c [22] and Pfs230D1M [38] on the surface of chimeric VLP. The small surface protein (dS) of the duck hepatitis B virus (DHBV) was used as VLP scaffold [41, 42]. The *P. falciparum* transmission stage antigens were genetically fused to the dS and the resulting fusion proteins were co-expressed with non-fused wild-type dS in recombinant strains of the methylotrophic yeast *Hansenula polymorpha* (*H. polymorpha*, syn. *Pichia angusta*, *Ogataea polymorpha*, [43]) which allowed the isolation of chimeric VLP composed of wild-type dS and the respective fusion protein. In contrast to previous VLP platforms, the dS-based VLP scaffold allows the stable incorporation of a variety of large molecular weight (MW) foreign antigens. In combination with the yeast expression system, this technology is highly productive and not limited to small scale fundamental research [44]. Thus, the key challenges in the field of chimeric VLP development are met [13,14,45] which makes this platform an attractive and competitive alternative to previously described VLP platforms [30–33]. Furthermore, expression of transmission-blocking antigens as VLP may enable the future co-formulation of these with RTS,S in multistage vaccines.

## 3. Materials and methods

### 3.1. Genes, plasmids and strains

Fusion proteins were designed by *N-*terminal fusion of malaria antigens to the VLP scaffold protein dS. Open reading frames (ORF) encoding the fusion proteins were synthesized by GeneArt/Life Technologies (Regensburg, Germany). They were codon-optimized for heterologous expression in *H. polymorpha* and flanked by *Eco*RI and *Bam*HI restriction sites. Synthesized ORF were inserted between the *Eco*RI and *Bam*HI sites of a derivative of the *H. polymorpha* expression plasmid pFPMT121 [46] which carried the *LEU2* instead *URA3* gene for selection in yeast. The sequences of the ORF post subcloning were confirmed by sequencing prior to yeast transformation. Cloning was done in bacterial strain *Escherichia coli* NEB^®^ 10-beta (New England Biolabs, Frankfurt a. M., Germany) grown at 37 °C in lysogeny broth [47] supplemented with 60 mg L^-1^ ampicillin (Applichem, Darmstadt Germany). The auxotrophic *H. polymorpha* strain ALU3 (relevant genotype: *ade1*, *leu2*, *ura3*) [48] derived from wild type strain ATCC^®^ 34438™ (CBS 4732, IFO 1476, JCM 3621, NBRC 1476, NCYC 1457, NRRL Y-5445) was used as expression host. Recombinant yeast cell lines were generated by electroporation [49] and a subsequent strain generation and isolation protocol [50]. Thereby, the expression plasmids integrated genomically stable in different copy numbers into the host genome. Heterologous yeast strains were stored as glycerol stocks at − 80 °C. Recombinant *H. polymorpha* strains co-producing the dS and a fusion protein were generated by the “staggered transformation approach” and screened as previously described [44].

### 3.2. Yeast cell mass generation

#### 3.2.1. Shake flask

VLP composed of Pfs230D1M-dS and dS were purified from cell mass of strain Ko#119, grown in 2 L baffled shake flasks filled with 200 mL YPG medium containing 20 g L^−1^ glycerol (AppliChem, Darmstadt, Germany) as carbon source and 0.1 g L^-1^ adenine (AppliChem, Darmstadt, Germany). A pre-culture grown in YPD medium to stationary phase was used as inoculum. The main cultures were incubated at 37 °C and 130 rpm with 5 cm throw. After 56 h of derepression, 1 % (v/v) methanol was added to the cultures for induction of target gene expression. After 72 h total cultivation time, cells were harvested by centrifugation (6,000*g*, 15 min, 4 °C), washed once with wash buffer (50 mM Na-phosphate buffer, 2 mM EDTA, pH 8.0) and stored at −20 °C.

#### 3.2.2. Bioreactor

VLP containing the fusion proteins Pfs25-dS or Pfs230c-dS were purified from cell mass of strain RK#097 or RK#114, respectively. Strains were grown in a 2.5 L scale stirred tank bioreactor (Labfors 5, Infors, Bottmingen, Switzerland). It was sterilized by autoclaving after filling with 2.5 L animal component free complex medium containing 20 g L^-1^ yeast extract (BD Biosciences, Heidelberg, Germany), 40 g L^-1^ peptone from soymeal (Applichem, Darmstadt Germany), 10 g L^-1^ glycerol, 11 g L^-1^ glucose-monohydrate, and 0.1 g L^-1^ adenine. Aqueous solutions of NH_3_ (12.5 % (w/w), sterile filtered) and H_3_PO_4_ (28 % (w/w), Merck, Darmstadt, Germany) were used as corrective media to keep pH constant (set point 6.5) throughout fermentation and Struktol J 673 (10 % (v/v) aqueous solution, Schill+Seilacher, Hamburg, Germany) was utilized as antifoam agent. Aeration was adjusted to 1 vvm (2.5 NL min^-1^) and the medium was inoculated to an optical density (OD_600_) of 0.6 using shake flask pre-cultures.

After a batch phase of 8 h, strain RK#114 was fed continuously with 275 mL of derepression solution (750 g L^-1^ glycerol) over 31 h. Formation of product was then induced by pulse-wise addition of 100 mL induction solution (285 g L^-1^ glycerol and 715 g L^-1^ methanol). Cells were harvested by centrifugation (6000*g*, 15 min, 4 °C) after 72 h total cultivation time, washed with wash buffer (25 mM Na-phosphate buffer, 2 mM EDTA, pH 8.0) and stored at −20 °C until further processing.

Strain RK#097 was fed with 160 mL of derepression solution over 36.5 h after the batch phase. 90 mL induction solution were added pulse-wise and fermentation was stopped after 65.2 h total cultivation time. Cells were harvested and stored as described before.

The dry cell weight (DCW) was quantified using a moisture analyzer (MLS 50-3 HA250, Kern & Sohn, Balingen, Germany). OD_600_ of cell suspensions was determined with a spectrophotometer (DU 640 Beckman Coulter, Brea, California, USA).

### 3.3. Purification of VLP

All VLP preparations were formulated in desalting buffer (8 mM Na-phosphate buffer pH 7, 154 mM NaCl) at concentrations in the range of mg mL^-1^. However, the purification protocols steps were adjusted for the different chimeric VLP.

#### 3.3.1. Purification of Pfs25-dS/dS VLP

Pfs25-dS/dS VLP were purified from strain RK#097 in preparative manner as described before [44, 51]. Briefly, cells were disrupted by six cycles of high pressure homogenization (∼1500 bar, APV 2000, SPX Flow Technology, Unna, Germany) and the cell homogenate was adjusted to 4.5 % (w/w) PEG_6000_ and 0.45 M NaCl. After incubation over-night at 4 °C and subsequent centrifugation (17,000*g*, 30 min, 4 °C), the product was adsorbed to fumed silica matrix Aerosil (type 380 V, Evonik, Essen, Germany). The matrix was washed once with 77 mM NaCl aqueous solution. Desorption buffer (10 mM di-sodium tetraborate decahydrate, 2 mM EDTA, 6 mM deoxycholic acid sodium salt, pH 9.1) was used to remove the product from the Aerosil (1 h, 25 °C). The desorbed material was applied to ion exchange chromatography (Mustang Q XT, PALL Life Sciences, Port Washington, New York, United States). Product containing fractions were pooled and concentrated by ultrafiltration (Minimate™ TFF tangential flow filtration Capsule Omega 100 k Membrane, PALL, Port Washington, New York, United States) prior to CsCl density gradient ultracentrifugation (1.5 M CsCl) in Optima™ L90K centrifuge (rotor type: 70.1 Ti, tubes: 16 * 76 mm, Beckman Coulter, Brea, California, USA) for 65 h at 48,400 rpm and 4 °C. Product containing fractions were pooled, desalted by dialysis (Slyde-A-Lyzer™ dialysis cassettes, MWCO 20 kDa, Thermo Fisher Scientific, Waltham, USA) against desalting buffer (8 mM Na-phosphate buffer pH 7, 154 mM NaCl, AppliChem, Darmstadt, Germany) and 0.45 µm filtered (Filtropur S 0.45 filters, Sarstedt, Nümbrecht, Germany).

#### 3.3.2. Purification of Pfs230c-dS/dS VLP

The following adjustments were made for the purification of Pfs230c-dS/dS VLP from strain RK#114: 2.5 % (w/w) PEG_6000_ and 0.25 M NaCl were used for clarification of the crude cell lysate after high pressure homogenization. Following the Aerosil batch adsorption step, the silica matrix was washed twice applying aqueous solution of 77 mM NaCl and 2.5 mM deoxycholic acid sodium salt or aqueous solution of 77 mM NaCl. The desorbed material was then subjected two consecutive times to Capto Core 700 chromatography matrix (GE Healthcare, Amersham, UK) applying 5 mg protein per ml resin. The unbound product fraction was concentrated by ultrafiltration and the retentate was applied to CsCl density gradient ultracentrifugation as described for the Pfs25-dS/dS VLP purification. Product containing fractions were then pooled and dialyzed against desalting buffer in two steps: first, the CsCl concentration was reduced to 0.5 M CsCl. Then, 0.05 % (w/v) SDS were added before the dialysis against desalting buffer was continued. The dialyzed sample was 0.45 µm filtered.

#### 3.3.3. Purification of Pfs230D1M-dS/dS VLP

The protocol for purifying Pfs230D1M-dS/dS VLP from strain Ko#119 was modified compared to the purification of Pfs230c-dS/dS VLP from strain RK#114. A 100 mM Na-carbonate/bicarbonate buffer (pH 9.2) with 1.2 M urea [52] was used for desorption of the product from the fumed silica matrix Aerosil. The desorbate was subjected to only one run of Capto Core 700 chromatography. Ultrafiltration, CsCl density gradient ultracentrifugation, dialysis and filtration were performed as described for Pfs25-dS/dS VLP purification from strain RK#097.

### 3.4. Protein, lipid and VLP analysis

Protein concentrations were determined with the Pierce^TM^ BCA protein Assay kit (Thermo Fisher Scientific, Waltham). Lipid content of VLP preparations was determined based on sulfo-phospho-vanillin reaction [53] with refined soya oil (Caesar & Loretz GmbH, Hilden, Germany) used as standard.

Sodium dodecyl sulfate polyacrylamide gel electrophoresis (SDS-PAGE) and Western blotting were performed as previously described [44]. In short: The Criterion^TM^ system from BioRad (München, Germany) was used to run SDS-PAGE. Cellulose nitrate membranes (Sartorius Stedim Biotech, Göttingen, Germany) were used for semi dry Western blotting of the proteins. Membranes were subsequently blocked by 3 % powdered milk in PBS containing 0.05 % Tween 20 (Pfs230-related Western blots) or Roti^®^-Block (Carl Roth GmbH, Karlsruhe, Germany, Pfs25-related Western blots). Primary antibodies used for immunolabelling (Table *1*) were utilized in combination with appropriate secondary antibodies purchased from BioRad (München, Germany) and BCIP-NBT (VWR international, Radnor, USA) or HRP substrate (Thermo Fisher Scientific, Waltham, USA).

**Table 1.** List of immunoreagents used for specific detection of the target proteins.

| Antigen | Primary antibody | Source | Reference |
| --- | --- | --- | --- |
| dS | 7C12<sup>(a)</sup> | BioGenes GmbH (Berlin, Germany) | [56,57] |
| Pfs25 | 32F81<sup>(b)</sup> | National Institutes of Health (NIH, Bethesda, Maryland, USA) | [9,58] |
|  | 4B7<sup>(b)</sup> | National Institutes of Health (NIH, Bethesda, Maryland, USA) | [23,59] |
| Pfs230 | mouse polyclonal<sup>(c)</sup> | National Institutes of Health (NIH, Bethesda, Maryland, USA) | [55] |
|  | 1B3 | National Institutes of Health (NIH, Bethesda, Maryland, USA) | [10] |

(a) detects the wild-type dS and the dS domain of each fusion protein
(b) Monoclonal antibodies 32F81 and 4B7 were kindly provided by PATH Malaria Vaccine Initiative and BEI Resources NIAID, NIH.
(c) Polyclonal mouse antibody was kindly provided by Carole Long and Kazutoyo Miura, NIH.

Coomassie staining of polyacrylamide (PAA) gels were also done as previously described [54]*. N*-Glycosylation of the heterologous target proteins was analyzed by treatment with an endoglycosidase H (EndoH) prior to SDS-PAGE [44].

Host cell proteins (HCP) were quantified by anti-HCP enzyme-linked immunosorbent assay (ELISA) as described previously [44].

Analyses by ELISA were performed in Nunc MaxiSorp™ flat-bottom 96 well ELISA plate (Thermo Fisher Scientific, Waltham). For specific detection of *P. falciparum* antigen Pfs25, the wells were coated over-night at 4 °C with 1 µg mL^-1^ (50 µL per well) of the respective VLP in PBS and blocked with 1 % (w/v) BSA in PBS for 1 - 2 h at RT before polyclonal mouse anti-Pfs25 antibodies were applied as primary immunoreagents for 2 h. The plate was washed thrice using PBST in between antibody incubation steps. Secondary polyclonal goat anti-mouse IgG HRP-conjugated antibody was used to detect antibody binding. Color detection was developed using ABTS liquid substrate (Sigma-Aldrich) which was subsequently stopped with 1 % SDS. The level of antibody binding was measured as optical density in a GENios Microplate Reader (Tecan, Männedorf, Switzerland) at 405 nm.

For specific detection of *P. falciparum* antigen Pfs230, plates were coated with indicated concentrations of VLP (4°C, over-night) and subsequently blocked with 1% casein in PBS (Sigma-Aldrich) for 2h at 37°C before primary antibodies were added (polyclonal mouse anti-Pfs230 or monoclonal 1B3 antibody, 10 µg mL^-1^). Secondary HRP-conjugated antibodies (polyclonal goat anti-mouse IgG at 1/1000 from Millipore) were used to detect antibody binding. Color detection was developed using ABTS liquid substrate (Sigma-Aldrich), which was subsequently stopped using 1 % SDS. PBS was used as a negative control and plates were washed thrice using PBS with 0.05 % Tween in between antibody incubation steps. The level of antibody binding was measured as absorption at 405nm (A_405nm_).

Analysis of VLP were performed as essentially described previously described [44] by dynamic light scattering (DLS), super-resolution microscopy (N-SIM; structured illumination microscopy) and transmission electron microscopy (TEM). As primary anti-Pfs230 antibodies, polyclonal mouse antibodies were applied that were described to have transmission-blocking activity [55]. Cross-reactivity with plain dS VLP without a fusion protein was checked carefully.

## 4. Results

### 4.1. Design of fusion proteins

In this study, three different types of chimeric VLP each displaying a different foreign antigen derived from *P. falciparum* were developed. The incorporation of *P. falciparum* transmission stage antigens into the dS-based VLP scaffold was realized by the design of fusion proteins each containing one of the malaria antigens *N-*terminally fused to the dS. The formation of VLP was allowed by interaction of dS subunits or dS domains of fusion proteins. Literature was screened for promising targets and the following antigens were chosen to be displayed on the surface of the membranous VLP. Additional details on the fusion protein construction are given in Table 2.

**Table 2.** Summary of fusion proteins constructed and recombinantly produced in *H. polymorpha*.

| Fusion protein | Number<br>of aa<sup>(a)</sup> | MW<br>[kDa] | Genbank<br>reference of CDS | *P. falciparum* donor protein<br>(Genbank reference) | Donor protein domain<br>fused to dS | aa linker between *P.*<br>*falciparum* antigen and dS |
| --- | --- | --- | --- | --- | --- | --- |
| Pfs25-dS | 341 | 37.0 | MH142260 | zygote/ookinete surface protein<br>Pfs25 (XP_001347587.1) | aa 23-193 [25] | GAGA |
| Pfs230c-dS | 861 | 98.0 | MH142261 | gametocyte surface protein<br>Pfs230 (XP_001349600.1) | aa 443-1132 [22] | GAGA |
| Pfs230D1M-dS | 366 | 40.3 | MH142262 | gametocyte surface protein<br>Pfs230 (XP_001349600.1) | aa 542-736<sup>(b)</sup> [38] | GAGA |

<sup>(a)</sup> Including an artificial start-methionine and aa 2-167 of the dS at the C-terminus
<sup>(b)</sup> Including N585Q aa exchange, elimination of a potential *N*-glycosylation site

The fusion protein Pfs25-dS comprised amino acids (aa) 23-193 of the cysteine-rich zygote/ookinete surface protein Pfs25 of *P. falciparum* fused to the dS. The Pfs25 part contained the four EGF-like domains of the naive antigen but missed the *N*-terminal signal sequence and the hydrophobic C-terminus [24,25,55]. Pfs230c-dS was constructed by fusing a 630 aa fragment (Pfs230c, [22]) of the *P. falciparum* transmission stage antigen Pfs230 to the dS. Due to its size, this fragment was challenging for a chimeric VLP-based approach. Thus, a shorter variant thereof (Pfs230D1M, aa 542-736 according to MacDonald et al. [38]) was introduced in the third fusion protein construct, Pfs230D1M-dS; this construct has been effectively expressed as a monomeric protein in *Pichia pastoris* and is a TBV candidate in clinical development [38]. All fusion protein encoding genes were inserted into a pFPMT121-based plasmid [46] which carried the *LEU2* gene for selection of transformed yeast strains.

### 4.2. Isolation of recombinant *H. polymorpha* production strains

Typically, co-production of dS and a fusion protein composed of a foreign antigen fused to dS allows formation of chimeric “antigen-dS/dS” VLP [41, 42]. For the generation of the three different types of VLP each displaying one of the *P. falciparum-*derived antigens, three recombinant *H. polymorpha* cell lines needed to be isolated that co-produce the scaffold protein dS and Pfs25-dS, Pfs230c-dS or Pfs230D1M-dS. To generate such strains, the dS-producing cell line A#299 [44] was super-transformed with an expression plasmid encoding the respective fusion protein. From each of the transformations, one strain co-producing the dS and the respective fusion protein was selected from the resulting transformants and used for production of chimeric VLP containing dS in combination with Pfs25-dS, Pfs230c-dS or Pfs230D1M-dS. The recombinant *H. polymorpha* strains that were used for production of VLP are indicated in Table 3.

**Table 3.** Recombinant *H. polymorpha* production strains used for the generation of VLP.

| Yeast strain designation | Transformed host strain | Produced recombinant protein(s) | Relevant genotype |
| --- | --- | --- | --- |
| RK#097 | A#299<sup>(a)</sup> | dS and Pfs25-dS | *URA3, LEU2, ade1* |
| RK#114 | A#299<sup>(a)</sup> | dS and Pfs230c-dS | *URA3, LEU2, ade1* |
| Ko#119 | A#299<sup>(a)</sup> | dS and Pfs230D1M-dS | *URA3, LEU2, ade1* |

<sup>(a)</sup> Recombinant dS-producing *H. polymorpha* strain previously described [44]

### 4.3. Production of chimeric Pfs25-dS/dS VLP

Chimeric Pfs25-dS/dS VLP composed of wild-type dS and the fusion protein Pfs25-dS were isolated from cell paste of strain RK#097. A total of 97.6±10.3 mg chimeric VLP could be isolated from 97±3 g DCW that were processed (1.0±0.1 mg g^-1^). In denaturing assays (SDS-PAGE, Western blot, Fig 1 A), the final sample was compared to plain dS VLP containing the dS without fusion protein [44] and thus lacks the fusion protein-specific signals. Apart from that, similar protein signal patterns were observed for the chimeric Pfs25-dS/dS VLP preparation in comparison to the plain dS VLP. Analysis of the Coomassie stained PAA gel by densitometry (lane 2) indicated 90 % Pfs25-dS/dS purity on the protein level and about 3 % fusion protein content. Anti-dS and anti-Pfs25 Western blots (lanes 4 and 6) were used to identify the VLP forming proteins. The apparent MW of Pfs25-dS (∼33 kDa) and dS (∼15 kDa) were slightly below their theoretical MW of 37 kDa or 18.2 kDa, respectively. For both VLP preparations additional signals were detected in the anti-dS Western blot that likely correspond to either oligomeric forms (dimers, trimers, etc.) of the dS or authentic forms of higher mobility (dS-HMF, [44]). The Pfs25-dS was reactive with transmission-blocking mAb 32F81 but non-specific cross reactivity with the dS was observed (lanes 5 and 6) applying Roti^®^-Block (Carl Roth GmbH) as blocking reagent. Cross reactivity was not observed if 3 % powdered milk in PBS containing 0.05 % Tween 20 was applied as blocking reagent (Fig S 1 in the supplementary material).

**Figure 1.**
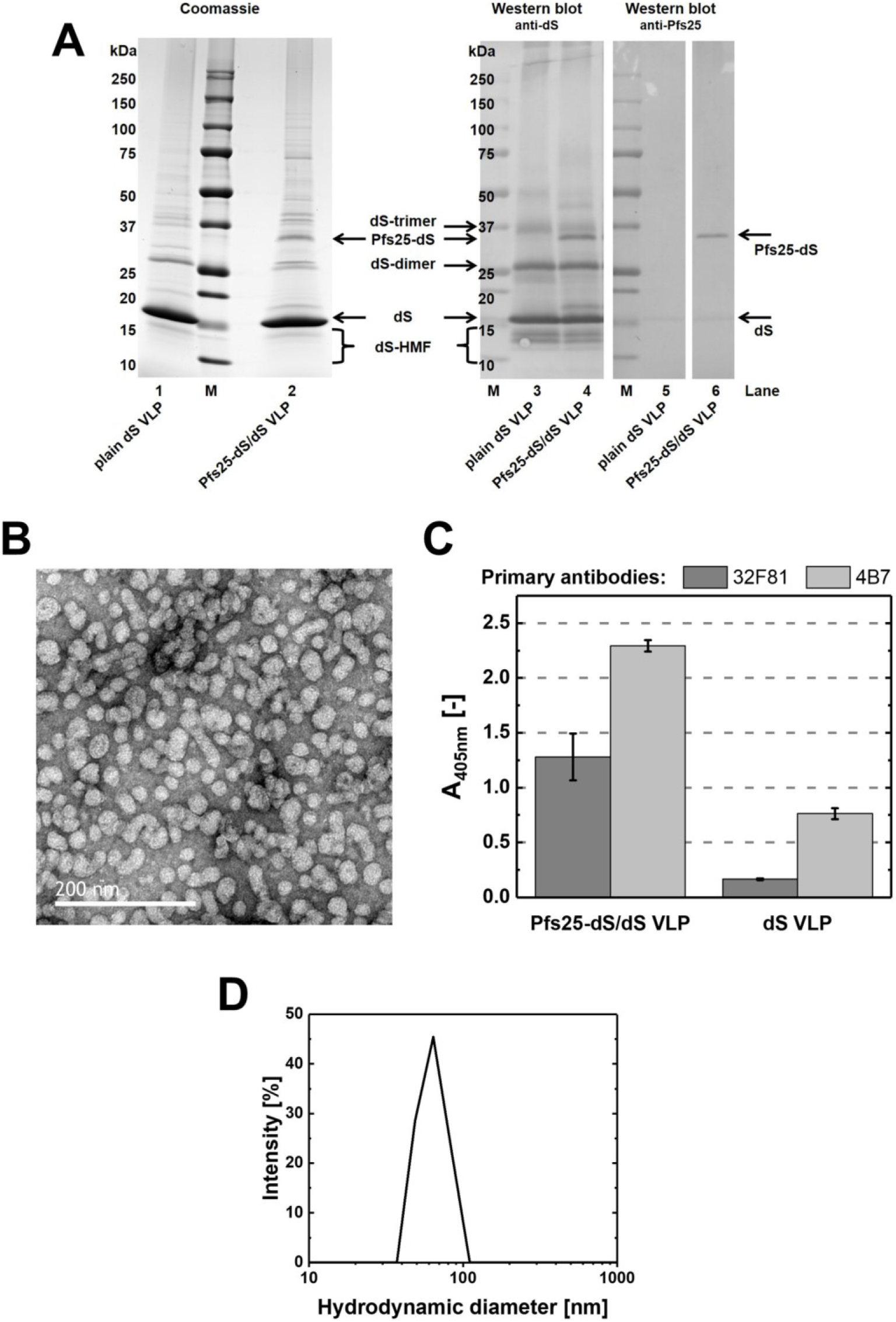
Analyses of purified Pfs25-dS/dS VLP derived from strain RK#097. (A): Reducing SDS-PAGE (left, 10 µg protein loaded) and Western blot (right, 1 µg protein loaded) comparing Pfs25-dS/dS VLP to plain dS VLP which were purified from strain A#299 and do not contain any fusion protein. Lanes 1 and 2: Coomassie stained PAA gel. Lanes 3 and 4: Western blot probed with anti-dS 7C12 mAb. Lanes 5 and 6: Western blot probed with anti-Pfs25 mAb 32F81 and analyzed on the same membrane. M: molecular weight marker. (B): TEM imaging. (C): Analysis by ELISA in comparison to plain dS VLP purified from strain A#299. The wells of the ELISA plate were coated with 1 µg mL^-1^ (50 µL per well) chimeric Pfs25-dS/dS VLP or same amounts of plain dS VLP. Error bars indicate standard deviation based on triplicates. (D): Size distribution determined by DLS.

ELISA was used to detect expression of Pfs25 on the surface of native VLP. Reactivity of the Pfs25-dS/dS VLP preparation was demonstrated with two Pfs25-specific antibodies 32F81 [9] and 4B7 [23] having transmission-blocking activity (Fig 1 C). Just as in the anti-Pfs25 Western blot, cross reactivity to the dS was observed. However, the Pfs25-dS/dS VLP were substantially more reactive.

Analysis by negative staining TEM and DLS (Fig 1 B and D) confirmed the formation of homogeneous particles. TEM imaging indicated particles of predominantly 20-40 nm according to manual evaluation. DLS showed a monomodal size distribution and a monodisperse particle population characterized by a hydrodynamic diameter of 64 nm (PDI 0.11). A summary of the production process and the composition of the final Pfs25-dS/dS VLP preparation can be found in Table 4.

**Table 4.** Summary of production processes leading to the three VLP preparations containing Pfs25-dS, Pfs230c-dS or Pfs230D1M-dS.

|  | VLP forming proteins |  |  |
| --- | --- | --- | --- |
|  | dS and Pfs25-dS | dS and Pfs230c-dS | dS and Pfs230D1M-dS |
| *H. polymorpha* strain | RK#097 | RK#114 | Ko#119 |
| Cell mass generation | Fermentation 2.5 L scale | Fermentation 2.5 L scale | Shake flask |
| DCW used for VLP purification [g] | 97±3 | 73.4±8 | 8.1±0.9 |
| VLP purity on protein level<sup>(a)</sup> [%] | 90 | 64 | 72 |
| Isolated VLP [mg] | 97.6±10.3 | 14.5±2.6 | 5.1±0.4 |
| VLP yield per biomass, Y<sub>P/X</sub> [mg g<sup>-1</sup>] | 1.0±0.1 | 0.2±0.06 | 0.64±0.1 |
| Product yield per culture volume [mg L<sup>-1</sup>] | 39±5 | 8±2 | 3.2±0.3 |
| Fusion protein per total VLP-forming protein<sup>(a)</sup> [%] | ~3 | ~30 | ~24 |
| VLP diameter by TEM [nm] | 20 - 40 | 44 - 60 | 42 - 62 |
| Hydrodynamic VLP diameter by DLS [nm] | 64 (PDI: 0.11) | 91 (PDI: 0.09) | 84 (PDI: 0.09) |
| Buoyant density [g cm<sup>-3</sup>] | 1.14 | 1.13 - 1.16 | 1.14 - 1.15 |
| Lipid content of purified VLP [%] | 30±4 | NE | 38±4 |

<sup>(a)</sup> Based on analysis of Coomassie-stained PAA gels by densitometry.
NE.: not evaluated

*N-*glycosylation of the Pfs25-dS fusion protein was analyzed in crude cell lysates by treatment with EndoH (Fig 2). The Pfs25-dS fusion protein construct “M” was described in Table 2 and was chosen for chimeric VLP production because its recombinant expression in *H. polymorpha* resulted in a homogeneous product (Fig 2, lanes 1 and 7). Despite its three potential *N*-glycosylation sites within the Pfs25 aa sequence, the “M” construct was not sensitive to EndoH treatment. The fusion protein was detected at ∼33 kDa MW without and after EndoH treatment indicating it was not *N-*glycosylated. However, in the initial experiments two additional Pfs25-dS constructs were included (“CL” and “QQ”, Fig 2). The “CL” construct analyzed in lanes 3/4 and 9/10 contained the leader sequence of the chicken lysozyme at its *N*-terminus instead of the artificial start-methionine of the “M” construct. The third construct (“QQ”) analyzed in lanes 5/6 and 11/12 was like the “CL” construct but contained two single amino acid exchanges (N112Q, N187Q) eliminating two of the three potential *N*-glycosylation sites. The design of the Pfs25-dS fusion protein affected its degree of *N*-glycosylation. The expression of “QQ” construct led to two Pfs25-dS species detected at ∼33 or ∼35 kDa MW (lanes 5 and 11). Expression of the “CL” variant resulted in the detection of four fusion protein species characterized by molecular weights of ∼33 kDa (faint band), ∼35 kDa, ∼38 kDa and ∼42 kDa (faint band, lanes 3 and 9). In both cases the signals unified in the ∼33 kDa signal after deglycosylation by treatment with EndoH (lanes 4, 6, 10 and 12). Thus the ∼33 kDa signal refers to the non-glycosylated form of the Pfs25-dS fusion protein.

**Figure 2.**
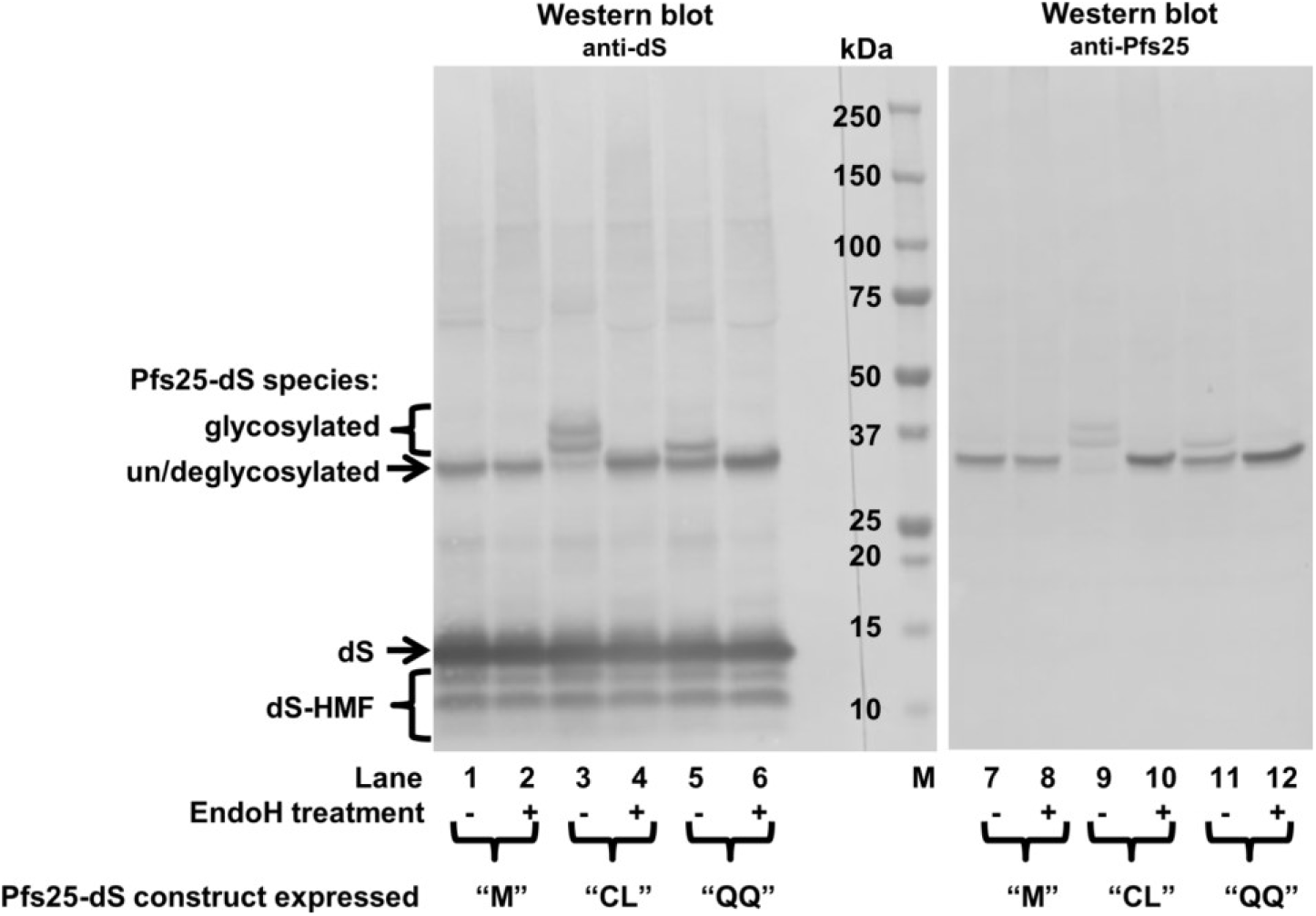
Western blot analysis of different Pfs25-dS constructs in crude cell lysates before and after treatment with EndoH. Crude cell lysates of three different recombinant *H. polymorpha* strains were analyzed by anti-dS (mAb 7C12) and anti-Pfs25 (mAb 32F81) Western blots. The strains co-expressed the wildtype dS and one of the three Pfs25-dS fusion protein constructs: “M” (*N-*terminal artificial start-methionine), “CL” (*N-*terminal chicken lysozyme signal sequence) or “QQ” (*N-*terminal chicken lysozyme signal sequence and N112Q, N187Q aa exchanges). Samples were treated with EndoH where indicated. M: molecular weight marker.

### 4.4. Production of chimeric Pfs230c-dS/dS VLP

From 73.4±8 g DCW of strain RK#114, 14.5±2.6 mg chimeric Pfs230c-dS/dS VLP composed of wild-type dS and the fusion protein Pfs230c-dS were isolated (Y_P/X_ = 0.2±0.06 mg g^-1^). At different stages during processing of Pfs230c-dS/dS VLP substantial losses of product were observed due to precipitation. Therefore, the purification protocol was adjusted compared to the purification of Pfs25-dS/dS VLP: the PEG and NaCl concentrations were reduced for clarification of the crude cell lysate, the Capto Core 700 matrix was used instead of the Mustang Q membrane adsorber and the dialysis procedure was modified. To reach higher product purity with the modified process, an additional wash step during the Aerosil batch procedure and a second Capto Core 700 run were added to the purification process.

Purified Pfs230c-dS/dS VLP were analyzed in native and non-native assays (Fig 3). VLP forming proteins were identified by anti-dS Western blot (Fig 3 A, lane 1) or by Pfs230-specific Western blot (lane 2), respectively. No cross-reactivity of the anti-Pfs230 polyclonal antibodies with the dS was observed. The purity on protein level (64 %) was investigated by densitometry for lane 3 of the Coomassie stained PAA gel (Fig 3 A). All bands detected between the fusion protein and the dS were considered as impurities. A subset of these bands was reactive in an anti HCP Western blot (Fig S 5 in the supplementary material). The most prominent host cell protein band beside the two product proteins (dS and Pfs230c-dS) was detected at 32 kDa apparent MW and represented 12 % of the total band volume of Coomassie stained lane 3. The Pfs230c-dS specific signal appears diffuse in lanes 3 and 4 (Fig 3 A). Upon treatment with EndoH, the diffuse smear disappeared and the main band became intensified by factor 2.6 according to analysis by densitometry. This revealed that the six potential *N*-glycosylation sites within the aa sequence of the Pfs230c antigen were in a subpopulation of the molecules at least partially occupied. Signal intensities of wild-type dS and deglycosylated Pfs230c-dS obtained by densitometry from lanes 5 to 8 were used to calculate the ratio of the two target proteins. For that purpose, the intensities were plotted against the protein amount loaded in the corresponding lane. The slopes obtained from linear regression revealed a ratio of wild-type dS to Pfs230c-dS in the chimeric Pfs230c-dS/dS VLP of approximately 70 % wild-type dS to 30 % fusion protein.

**Figure 3.**
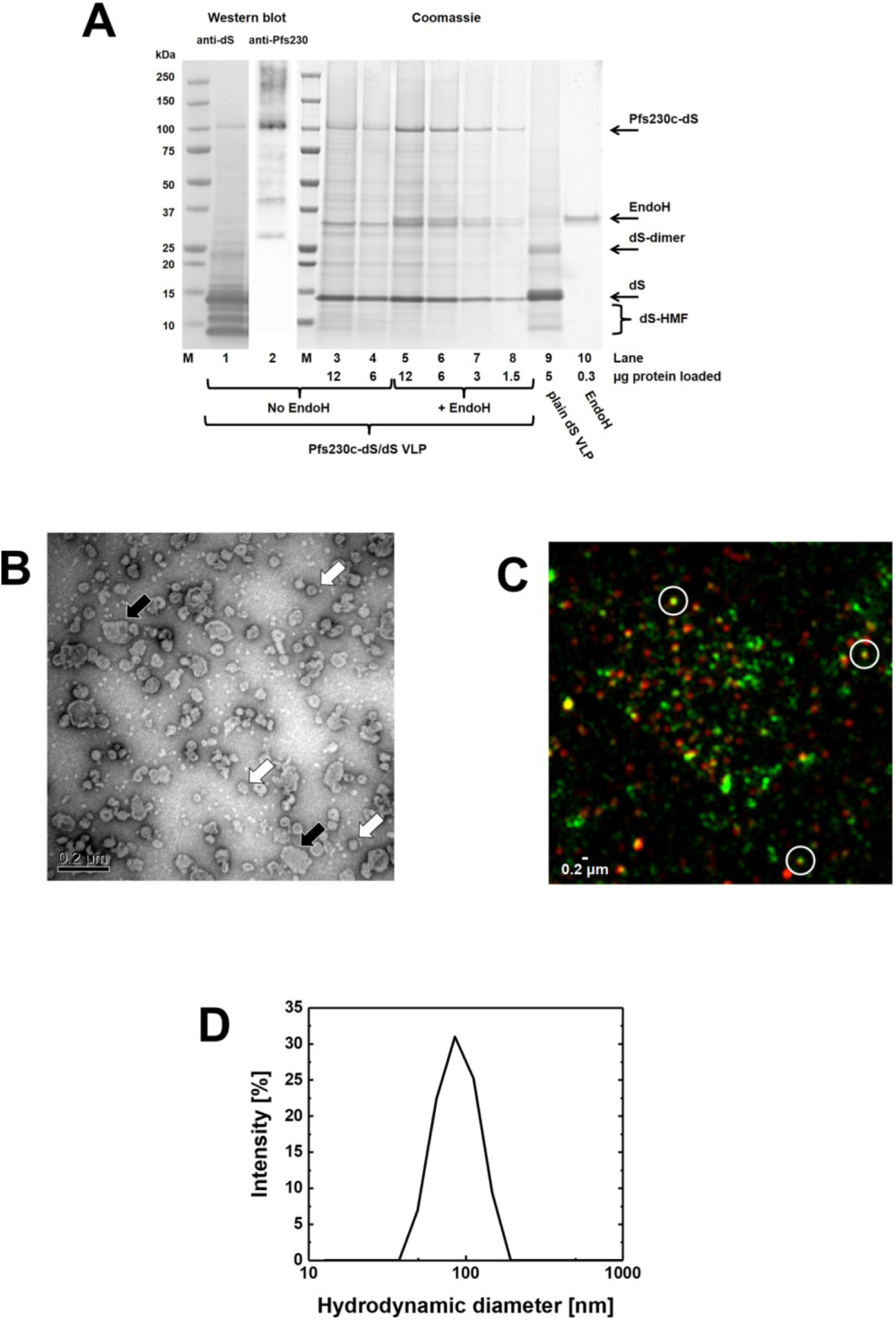
Analyses of purified Pfs230c-dS/dS VLP derived from strain RK#114. (A): Western blot probed with anti-dS 7C12 mAb (lane 1) or probed with polyclonal anti-Pfs230 antibody (lane 2) and Coomassie stained PAA gel (lanes 3-10). Samples loaded in lanes 5-8 were treated with an EndoH. Lane 9: purified plain dS VLP. Lane 10: EndoH used. M: molecular weight marker. (B): TEM imaging at 100,000-fold magnification. (C): N-SIM analysis of purified VLP containing Pfs230c antigen showing immunolabeling of dS (green) or Pfs230c (red). Three nano-scaled spots that showed co-localization of the fluorescence markers (yellow) were representatively circled. (D): Size distribution determined by DLS.

Formation of VLP was confirmed for both samples by TEM and DLS (Fig 3 B and D). DLS indicated a monomodal and monodisperse (PDI 0.09) particle population characterized by hydrodynamic diameter of 91 nm. However, the appearance of the Pfs230c-dS/dS VLP in TEM imaging was rather heterogeneous (Fig 3 B). The dominating species of detected objects were in the range of 44-60 nm diameter but also larger structures (>120 nm) were observed frequently and this could be due to particle aggregation.

Analysis by N-SIM was applied as native immunoassay using polyclonal antibodies described to have transmission-blocking activity [55]. This demonstrated co-localization of dS and Pfs230c antigens in nano-scaled particles (circled spots, Fig 3 C) providing supporting evidence of the formation of chimeric VLP by wild-type dS and Pfs230c-dS. Also, Pfs230c-dS/dS VLP were reactive with anti-Pfs230 polyclonal mouse antibodies [55] in ELISA (Fig 4 and Fig S 4) but failed to react with monoclonal anti-Pfs230 antibody 1B3 [10] (Fig S 4 in the supplementary material). Cross reactivity of the anti-Pfs230 polyclonal antibody with the dS VLP scaffold in form of plain dS VLP [44] was not observed in this native assay (Fig 4).

**Figure 4.**
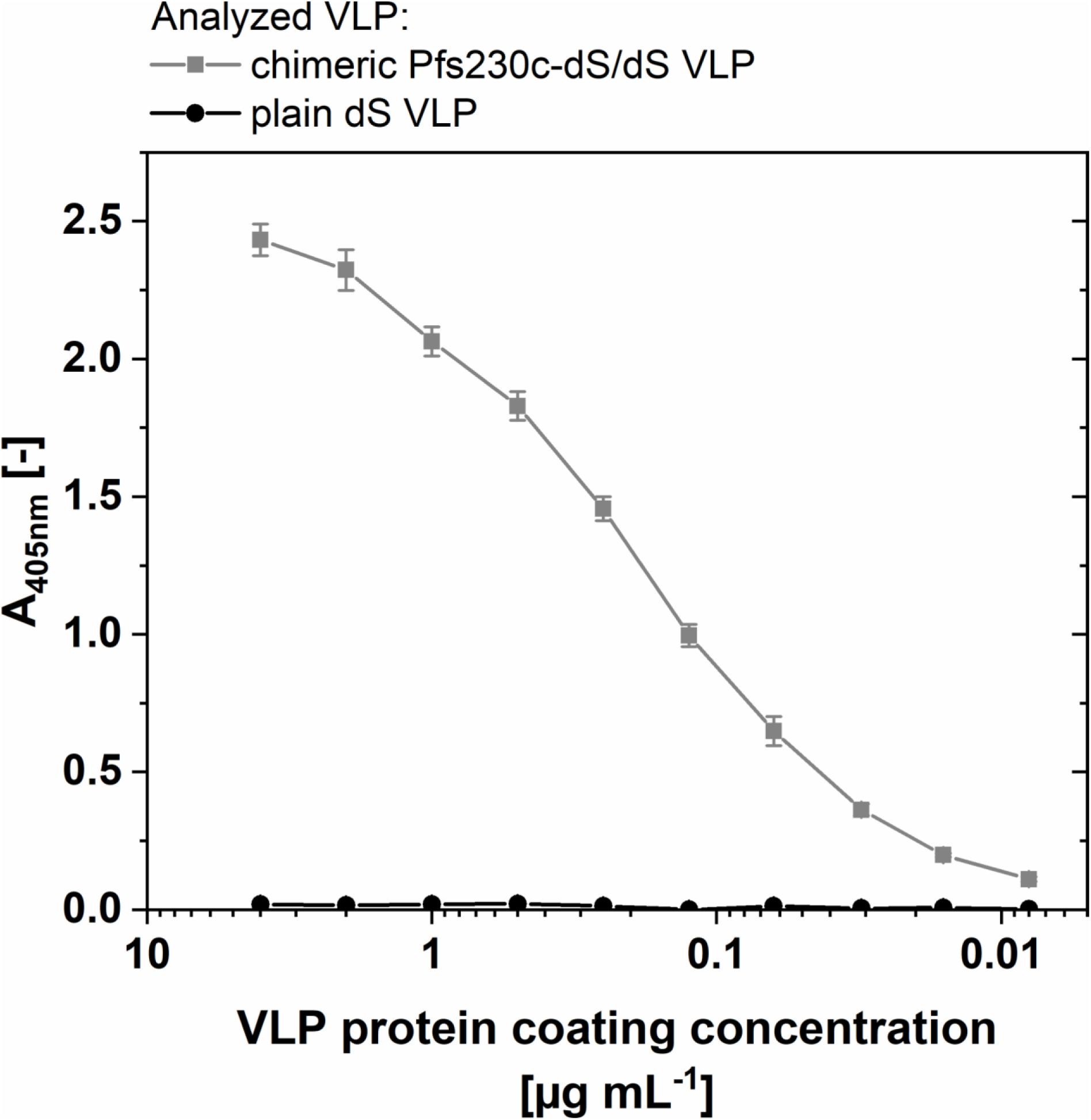
ELISA data on purified Pfs230c-dS/dS VLP derived from strain RK#114. The ELISA plate wells were coated with the indicated chimeric Pfs230c-dS/dS VLP or plain dS VLP. The mouse polyclonal antibody was applied at 10 µg mL^-1^.

### 4.5. Production of chimeric Pfs230D1M-dS VLP

Processing 8.1±0.9 g DCW of strain Ko#119 yielded 5.1±0.4 mg of chimeric Pfs230D1M-dS/dS VLP (Y_P/X_ = 0.64±0.1 mg g^-1^) that were composed of wild-type dS and the fusion protein Pfs230D1M-dS. Processing of Pfs230D1M-dS/dS VLP was easier than processing of Pfs230c-dS/dS VLP. No unexpected product losses during downstream processing (DSP) were observed and thus a less complex DSP could be chosen. The specific yield Y_P/X_ of Pfs230D1M-dS/dS VLP was about three times as much as for the Pfs230c-dS/dS VLP.

Both VLP-forming proteins were detected by anti-dS Western blot (Fig 5 A, lane 2). The fusion protein Pfs230D1M-dS was specifically detected by anti-Pfs230 Western blot (lane 3). Judging by their MW, the additional high MW signals in this lane likely correspond to oligomeric forms (dimers, trimers, etc.) of the fusion protein. Most likely, these forms were not detected by the anti-dS mAb in lane 2 because the signals were below the detection limit. Analysis of a Coomassie stained PAA gel (Fig 5 A, lane 1) by densitometry indicated 72 % purity on protein level and a composition of 24 % fusion protein and 76 % wild-type dS. Formation of VLP was confirmed by TEM (Fig 5 B) and indicated 42 – 62 nm diameter for the VLP. Size distribution analyzed by DLS (Fig 5 D) indicated monomodal size distribution and a monodisperse particle population characterized by a hydrodynamic diameter of 84 nm (PDI 0.09). N-SIM was performed as for Pfs230c-dS/dS VLP with a similar result (Fig 5 C). The Pfs230D1M-specific and the dS-specific signals co-localized in nano-scaled particles (circled spots).

**Figure 5.**
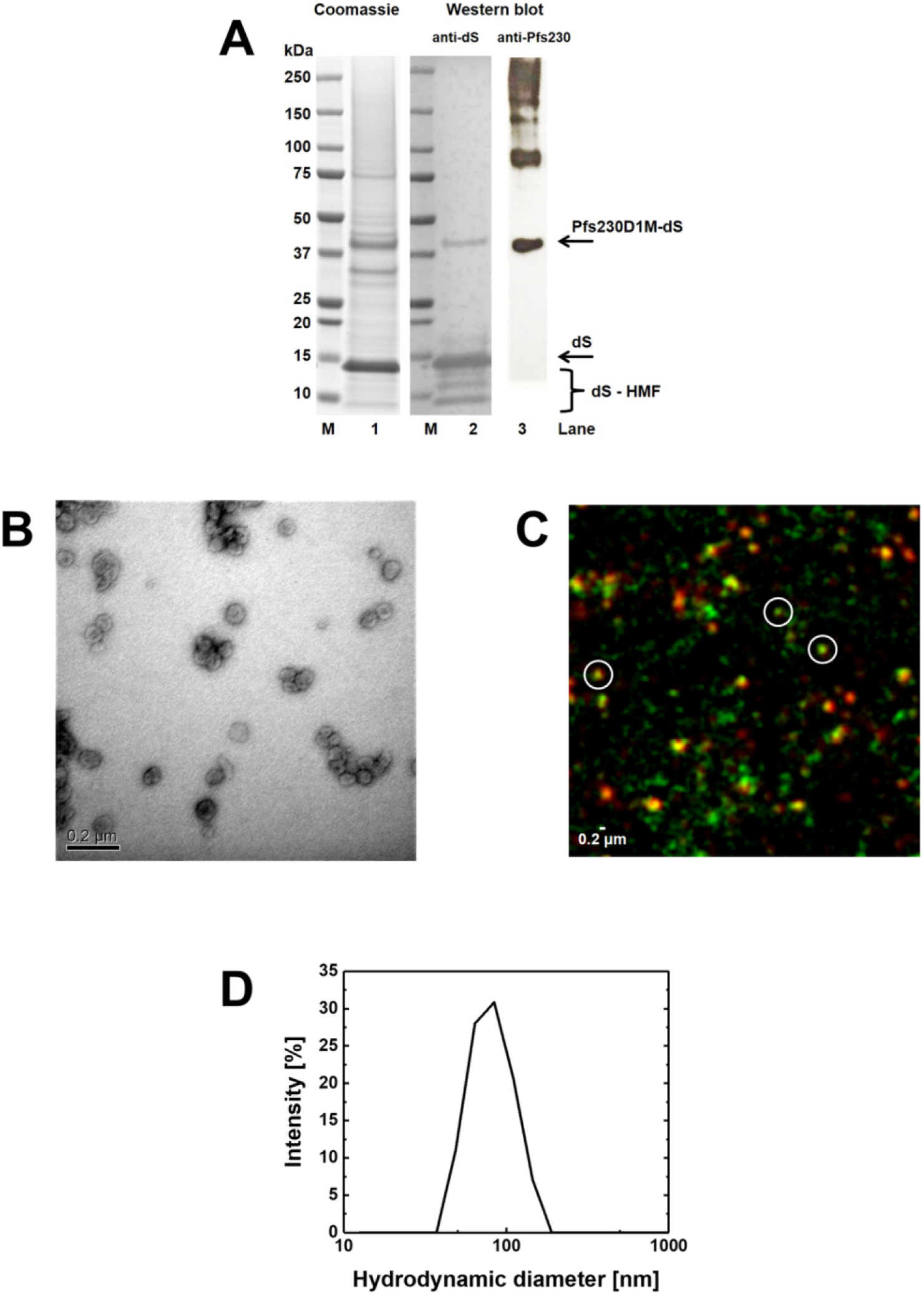
Analyses of purified Pfs230D1M-dS/dS VLP derived from strain Ko#119. (A): Coomassie stained PAA gel (lane 1, 12 µg protein loaded), Western blot probed with anti-dS 7C12 mAb (lane 2) or probed with polyclonal anti-Pfs230 antibody (lane 3). M: molecular weight marker. (B): TEM imaging at 100,000-fold magnification. (C): N-SIM analysis of purified VLP containing Pfs230D1M antigen showing immunolabeling of dS (green) or Pfs230D1M (red). Three nano-scaled spots that showed co-localization of the fluorescence markers (yellow) were representatively circled. (D): Size distribution determined by DLS.

### 4.6. Summary and comparison

The production processes and the compositions of the three different VLP preparations are summarized in Table 4. The purification of chimeric Pfs25-dS/dS VLP from strain RK#097 was the most productive process and yielded 1.0±0.1 mg VLP per g DCW or 39±5 mg VLP per L cell culture with 90 % purity on protein level. However, the fusion protein content in the VLP was lowest (∼3 %) in comparison to the other VLP preparations.

Despite the size of the Pfs230c antigen and difficulties due to product precipitation, isolation of chimeric Pfs230c-dS/dS VLP was successful. However, the VLP yield was considerably lower (0.2±0.06 mg g^-1^ or 8±2 mg L^-1^) than for the chimeric Pfs25-dS/dS VLP but the fusion protein content of the VLP was approximately 10-times higher than for Pfs25. The purity of Pfs230c-dS/dS VLP on the protein level was substantially lower (64 % on protein level) and could not be improved in the course of this study.

VLP yield and purity, were improved by modification of the fusion protein. The truncated Pfs230D1M-dS version led to tripling in VLP yield (Y_P/X_) combined with improvement in purity on the protein level by 8 %. However, the yield per culture volume was lower due to cultivation in shake flasks compared to fermentation in 2.5 L scale. The Pfs230D1M-dS/dS and the Pfs230c-dS/dS VLP preparations had comparable fusion protein contents.

The hydrodynamic diameters determined by DLS were consistently slightly larger than the respective diameters specified by manual evaluation of the TEM images. Nevertheless, all data collected are within dimensions that could be expected for this kind of VLP [44, 60]. The determined buoyant densities (1.13 - 1.16 g cm^-3^) and lipid contents (30-40 %) were also consistent throughout the VLP preparations and plausible for lipoproteins or VLP [61].

## 5. Discussion

This work introduces a novel VLP platform for the display of malaria transmission-blocking antigens derived from *P. falciparum* on the surface of a nano-scaled VLP scaffold. Chimeric VLP containing the leading malaria vaccine candidates Pfs25 and Pfs230 were engineered, purified and characterized.

A DSP approved for hepatitis B vaccine production from yeast [51] was used for isolation of highly pure chimeric Pfs25-dS/dS VLP. Initially, three different Pfs25-dS fusion protein constructs were constructed and it was shown that depending on the design of the construct, the degree of *N-*glycosylation varied (Fig 2). Due to product homogeneity and therewith potential regulatory advantages, the non-glycosylated variant was chosen for chimeric Pfs25-dS/dS VLP production. The purified Pfs25 containing VLP were reactive with transmission-blocking monoclonal antibodies 32F81 and 4B7 in ELISA [58, 59] and could be identified as particulate structures in TEM imaging (Fig 1). These can be considered as promising findings for upcoming immunization studies. However, cross reactivity of the dS with the antibodies 32D81 and 4B7 was observed in Western blot (Fig 1A) and ELISA (Fig 1C) which complicates interpretation of the results. In case of the antibody 4B7, the difference in reactivity with the chimeric Pfs25-dS/dS VLP or with plain dS VLP not containing the Pfs25 antigen was only about factor 3. This difference in reactivity could be expected to be greater and may be caused by misfolded Pfs25 antigen e.g. due to incomplete formation of disulfide bonds or because of the low Pfs25 fusion protein content in the isolated chimeric VLP. Depletion of Pfs25-dS relative to dS in the course of the DSP was not observed and thus this low inclusion rate could have two possible reasons.

1. The ratio of Pfs25-dS to wild-type dS produced by RK#097 was too low to facilitate isolation of chimeric VLP with higher fusion protein content.
2. The chosen fusion protein construct Pfs25-dS is suboptimal for high fusion protein content in this kind of VLP.

To overcome possibility 1, recombinant *H. polymorpha* strains producing improved fusion protein to dS ratios were generated by applying alternative strain generation approaches described in [44]. Solubility of the VLP forming proteins in homogenates of these new strains was however reduced compared to strain RK#097 (Figs S 2 and S 3 in the supplementary material). As a next step, solubilization of the VLP proteins would need to be optimized and purified chimeric VLP should then be compared head-to-head with purified particles from strain RK#097. To address possibility 2, the fusion protein construct may be modified e.g. by truncation of the Pfs25 antigen. The EGF2-domain of the protein was described to retain the transmission-blocking activity [62] and might be used instead of the more complex, full-length Pfs25.

For production of VLP expressing a Pfs230 construct, the 630 aa Pfs230c fragment [22] of the *Plasmodium* antigen Pfs230 was fused to the dS and chimeric Pfs230c-dS/dS VLP were purified. The incorporation of the 80 kDa Pfs230c fragment into chimeric VLP demonstrated that the integration of foreign antigens in dS-based VLP is not necessarily limited by their size; this is a substantial advantage over other VLP platforms [13]. The observed precipitation of the Pfs230c-dS containing material during DSP may be due to misfolded protein. This may also be an explanation for missing response in ELISA applying the 1B3 monoclonal antibody (Fig S 4 in the supplementary material). The Pfs230c-dS aa sequence contains 16 cysteine residues which could be linked incorrect via disulfide bonds. In our experiments, overexpression of a recombinant protein disulfide isomerase did not result in detectable reduction of product loss during DSP which does not support the hypothesis of incorrectly formed disulfide bonds. Additionally, our observations regarding the solubility of the Pfs230c construct correlate with reports on improving the solubility of Pfs230 fragments by choosing particular fusion partners [22, 63]. Although this issue did not hinder the isolation of chimeric VLP, it resulted in changes in the DSP and led to reduced VLP yields. Nonetheless, the obtained yields and fusion protein contents of the VLP (∼30 %) are good in the context of chimeric VLP vaccines [64] and the reactivity with the polyclonal anti-Pfs230 antibody in native ELISA (Fig 4) is a promising result.

A remaining challenge regarding the chimeric Pfs230c-dS/dS VLP purification is the relatively low purity of 64 % of the final preparation. It can be speculated that a contributor to the sub-optimal purity is that the residual HCP impurities were tightly associated with the particles or the Pfs230c antigen. Separation of product from contaminative proteins was not possible in the course of this study. A revised purification protocol may need to be developed addressing reduction of the most prominent, persisting protein contaminants already present in earlier purification steps.

Compared to the Pfs230c-dS/dS VLP, the Pfs230D1M [38] variant was easier to process; no loss of product due to precipitation was observed and higher VLP yield per biomass was achieved with comparable fusion protein content. This finding of improved solubility agrees with successful heterologous production of soluble Pfs230D1M fragment in *P. pastoris* and secretion of the product into the culture supernatant [38].

For the two chimeric VLP preparations containing Pfs230 fragments, the co-localization of dS and Pfs230 in nano-scaled particles was observed by N-SIM analysis. However, the resolution of the test set-up may not be sufficient for the detection of single chimeric VLP. Due to physicochemical homogeneity of the analyzed samples, it can be concluded that co-localization of both proteins in clusters of few VLP support the occurrence of both proteins in single VLP. Together, N-SIM analysis (Figs 3 C and 5 C) and ELISA (Fig 4) proved the accessibility of both Pfs230 constructs under native conditions for immunolabeling which substantiates the display of the respective malaria antigens on the VLP surface. Additionally, these native immunoassays demonstrated reactivity with antibodies that are known to have transmission-blocking activity. This is again promising regarding applying these VLP in vaccine immunogenicity studies.

The expression system *H. polymorpha* was shown to be a reliable and productive host for production of chimeric VLP displaying the difficult-to-express transmission-blocking antigens Pfs25 and Pfs230 on their surface. Yeasts combine the ease of genetic manipulation and the option for simple fermentation strategies of bacterial expression systems with the ability to modify proteins according to a general eukaryotic scheme [65]. Mammalian and insect cell expression systems might be the favorable systems in case of production and assembly of highly complex multi-layer VLP. However, the advantages of yeast-based VLP production is especially valued in the domain of simpler, single-layered VLP production [45,66–68]. Particularly, the methylotrophic yeast *H. polymorpha* should be considered for production of chimeric VLP vaccine candidates since it is already established as a safe and reliable microbial cell factory for the production of biopharmaceuticals like hepatitis B VLP vaccines [51,69,70] or recombinant products that have been granted “generally recognized as safe” (GRAS) status.

The generation of recombinant *H. polymorpha* strains is more laborious than for other yeast species, including *Saccharomyces cerevisiae* [50]. However, these additional difficulties are compensated by a number of positive characteristics which are advantageous in biotechnological applications (for review see e.g. [71–73]). These include mitotic stability of recombinant strains even under non-selective conditions due to stable integration of plasmids in high copy numbers into the host’s genome [50], the availability of strong and regulated promoters derived from the methanol utilization pathway [74], the applicability of different carbon sources, especially glycerol [75, 76] and the ease to grow *H. polymorpha* to high cell densities reaching dimensions of 100 g DCW per L culture broth [77]. Additional advantages of the methylotrophic yeast for the production of recombinant proteins are its relatively high optimal growth temperature of 37 °C which allows a better temperature management in large-scale fermentations and the tendency to reduced *N*-linked hyperglycosylation of recombinant proteins compared to *Saccharomyces cerevisiae* [74, 78] combined with the lack of the terminal, hyperallergenic α-1,3-linked mannose [79].

## 6. Conclusion

This study introduces a novel platform for the presentation of leading malaria transmission-blocking antigens of up to 80 kDa on the surface of chimeric VLP. Each of the generated chimeric VLP preparations was reactive under native conditions with antibodies described to have transmission-blocking activity. Regarding VLP yield, their purity and fusion protein content, the chimeric Pfs230D1M-dS/dS VLP appears to be the most promising candidate that emerged from this study. The obtained product yields in combination with the versatility and reliability of the described VLP production platform makes it a competitive system and should be considered for future malaria vaccine development. However, the potential of the three developed, chimeric VLP as effective vaccine candidates cannot be disclosed unless studies to assess their immunogenicity and transmission-blocking performance are completed. This represents together with improving the product purity especially for the chimeric Pfs230c-dS/dS VLP the future key tasks.

## Section S1: Additional data on the chimeric Pfs25-dS/dS VLP

### S1.1 Cross-reactivity of anti-Pfs25 antibodies

Fig 1 A of the main manuscript shows Western blot analyses on the chimeric Pfs25-dS/dS VLP (lanes 3 to 6). Cross reactivity of the anti-Pfs25 antibody with the dS scaffold protein was observed (Fig 1A, lanes 5 and 6). Therefore, Western blot analysis was repeated (shown in Fig S 1) with two different chimeric VLP preparations originating from two different *H. polymorpha* cell lines producing different amounts of two target proteins dS and Pfs25-dS. The blocking of the membrane was modified compared to the methodology described in the main manuscript. Instead of the Roti Block reagent (Carl Roth GmbH, Karlsruhe, Germany), 3 % (w/v) dry milk (in PBST) were applied as for the Pfs230-related Western blots. With this modified procedure we did not observe cross-reactivity of the anti-Pfs25 antibody with the dS in Western blot.

Lanes 2 and 4 in Fig S1 represent the actual Pfs25-dS/dS VLP preparation discussed in the main manuscript whereas lanes 1 and 3 represent a substantially lower concentrated Pfs25-dS/dS VLP preparation originating from a different *H. polymorpha* cell line. In both VLP preparation the dS was detected by the anti-dS antibody (lanes 1 and 2). However, only for the preparation applied to lanes 2 and 4 the presence of the Pfs25-dS fusion protein was substantiated by the anti-Pfs25 antibody. Potentially, the Pfs25-dS fusion protein content in the sample applied to lanes 1 and 3 is below the detection limit. However, cross reactivity with the VLP scaffold protein dS was not observed for neither of the samples most likely due to the altered Western blot procedure compared to the analysis shown in the main manuscript (Fig 1 A).

**Figure S 1.**
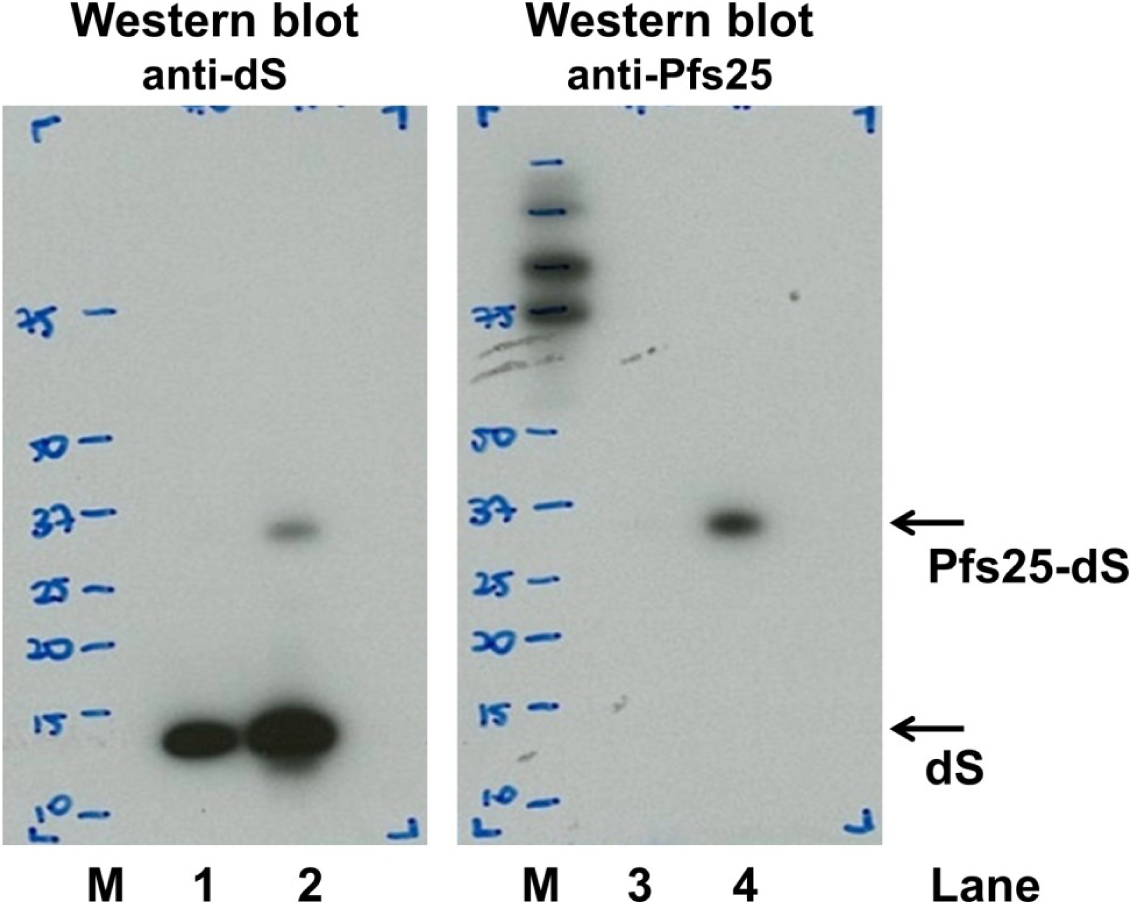
Repetition of Western blot analyses on chimeric Pfs25-dS/dS VLP. Chimeric Pfs25-dS/dS VLP preparations obtained from two different *H. polymorpha* cell lines were applied. Lanes 2 and 4: the Pfs25-dS/dS VLP preparation discussed in the main manuscript. Lanes 1 and 3: A lower concentrated VLP preparation derived from a different cell line expressing lower levels of dS and Pfs25-dS. Left: Membrane probed with anti-dS 7C12 mAb. Right: Membrane probed with anti-Pfs25 mAb 32F81 and analyzed on the same membrane. M: molecular weight marker

### S1.2 Reduced product solubilization at elevated Pfs25-dS expression levels

To overcome the low incorporation ratio of Pfs25-dS in the chimeric Pfs25-dS/dS VLP isolated from strain RK#097, additional strains co-expressing dS and Pfs25-dS were generated and screened for higher productive than strain RK#097 on the cell lysate level. One of them is the strain designated as DW#044. A side-by-side Western blot analysis of strain RK#097 and DW#044 is shown in Fig S 2. Cell pellets of the two strains were resuspended OD_600_ normalized in cell disruption buffer (25 mM Na-phosphate buffer, 2 mM EDTA, 0.5 % (w/v) Tween 20, pH 8.0). Cell disruption was carried out in 1.5 mL reaction tubes on a shaker (basic Vibrax® shaker, IKA®-Werke, Staufen, Germany) at maximal frequency for 30 min at 4 °C using glass beads (0.5–0.7 mm, Willy A. Bachofen, Nidderau-Heldenberg, Germany). One part of the resulting crude cell lysates was analyzed directly by anti-dS Western blot (lanes 1 and 4). The rest of the lysates was separated into soluble protein fraction (analyzed in lanes 3 and 6) and insoluble material (analyzed in lanes 2 and 5) by centrifugation (15 min, 13.000 *g,* 4 °C). The insoluble material was resuspended in distilled water volume-normalized to the volume of the centrifuged cell lysate prior to Western blot analysis. The comparison of lane 1 to lane 4 indicates higher productivity of the strain DW#044 compared to strain RK#097 regarding the fusion protein Pfs25-dS. However, in contrast to the material obtained from strain RK#097, the majority of the product proteins (dS and Pfs25-dS) produced by strain DW#044 was detected in the insoluble material (lane 5). Only a minority of the product was found in the soluble protein fraction (lane 6). The higher productivity on the cell lysate level (compare lanes 1 and 3) did not lead to higher product yields in the soluble protein fraction (compare lanes 3 and 6).

**Figure S 2.**
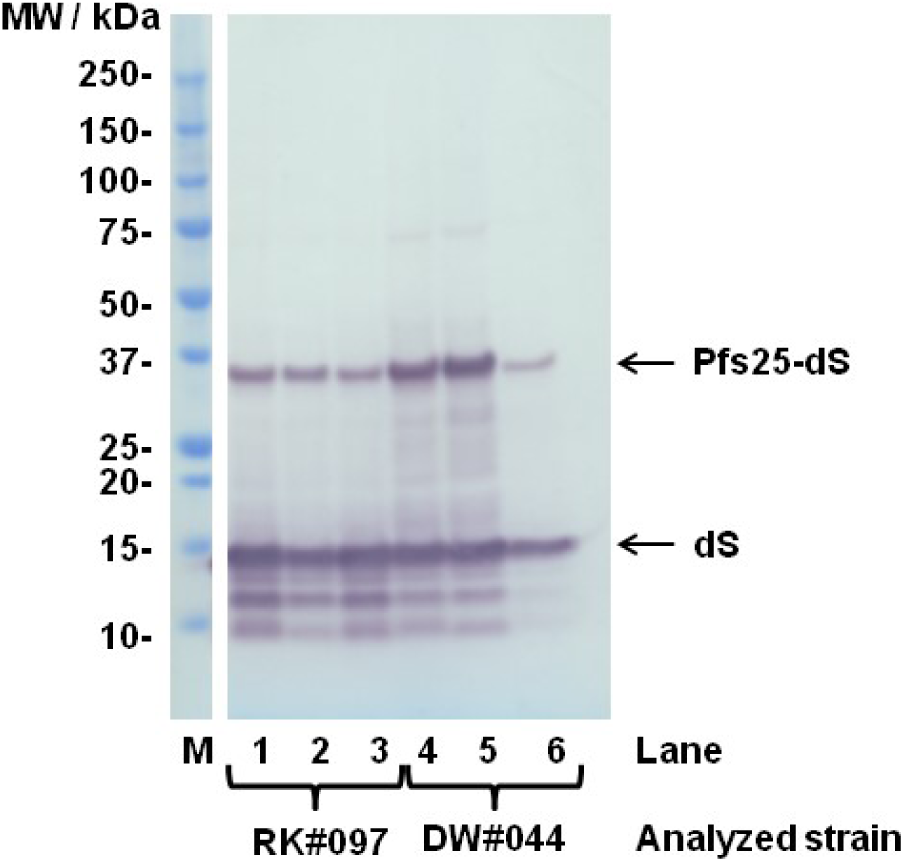
Side-by-side Western blot analysis of strains RK#097 and DW#044 co-producing the dS and Pfs25-dS. Cell lysates (lanes 1 and 4), insoluble materials (lanes 2 and 5) and soluble protein fractions (lanes 3 and 6) obtained from equal amounts of cells were applied to the gel. The membrane was probed with anti-dS mAb 7C12. M: molecular weight marker.

This was analyzed in more detail by anti-dS Western blot analyses applying dilution series of the crude cell lysates and the soluble protein fractions (Fig S 3). The methodology of Western blot is only a semi-quantitative approach and the results have to be treated with caution. However, the decreased solubilization of the target proteins in case of the strain DW#044 (Fig S 3 B) is obvious compared to the strain RK#097 (Fig S 3 A). Based on analysis by densitometry approximately 54 % of the target proteins are solubilized in case of the strain RK#097 whereas only ∼20 % of the target proteins were solubilized in case of the strain DW#044.

**Figure S 3.**
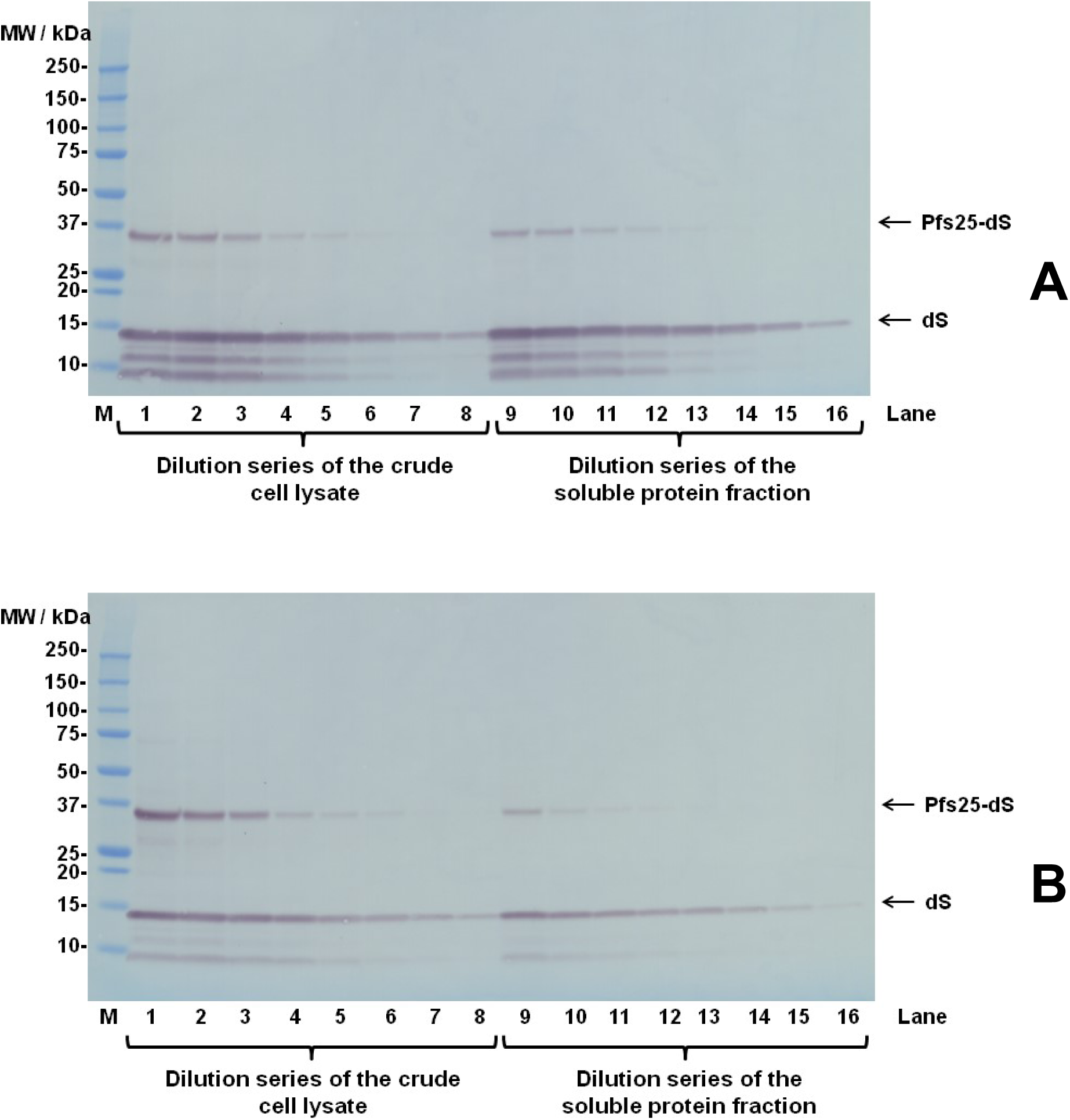
Western blot analyses of crude cell lysates and soluble protein fractions of strains RK#097 (A) and DW#044 (B). The fractions were applied as dilution series (factor 2 steps). The membrane was probed with anti-dS mAb 7C12. M: molecular weight marker.

## Section S2: Additional data on the chimeric Pfs230c-dS/dS VLP

### S2.1 Reactivity of different anti-Pfs230 immunoreagents with Pfs230c-dS/dS VLP in ELISA

During the development of methodologies to analyze the Pfs230c-dS/dS VLP, different primary immunoreagents were tested. The mouse polyclonal antibody was found to be substantially more reactive than the 1B3 monoclonal antibody (Fig S 4).

**Figure S 4.**
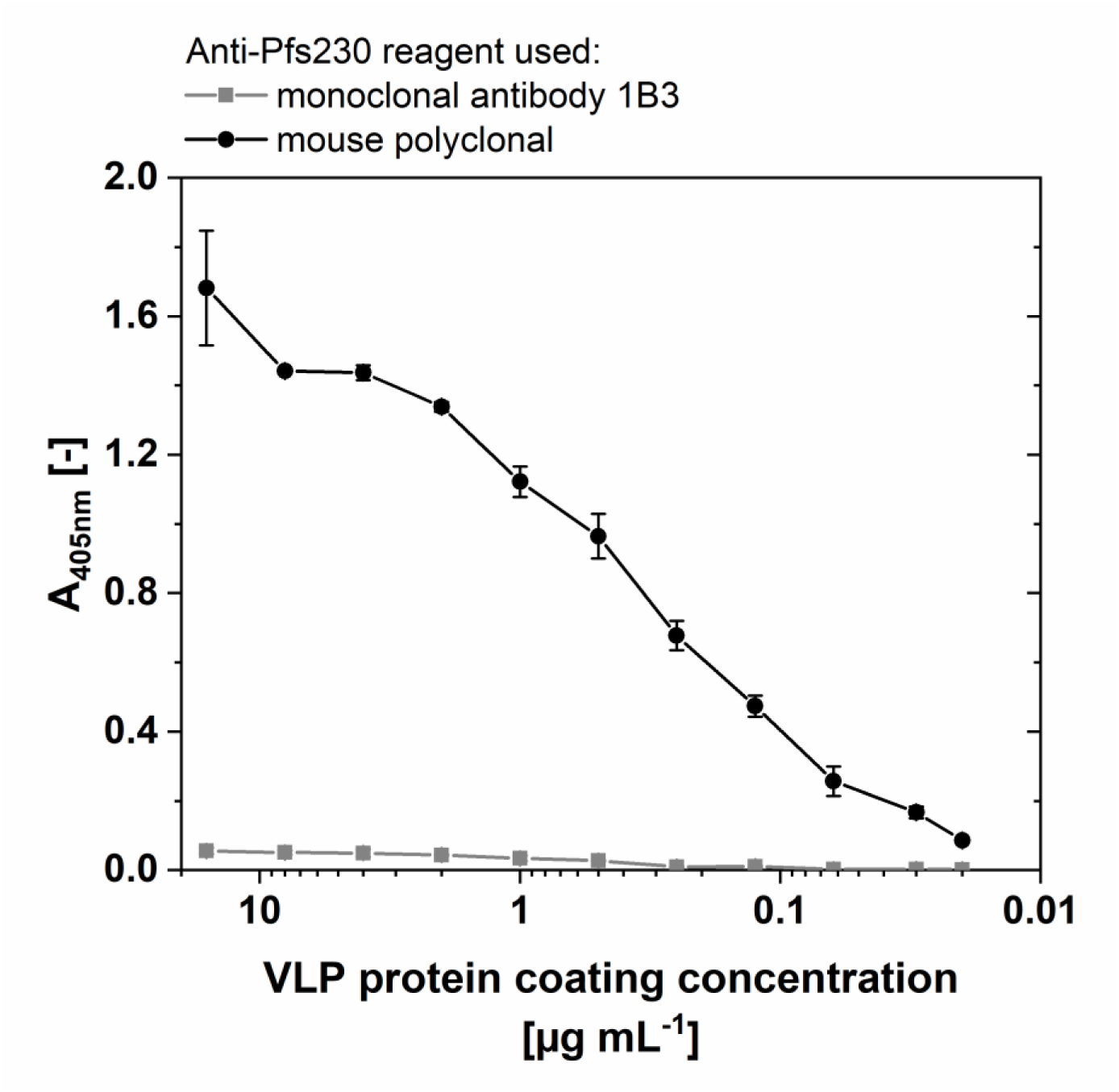
ELISA data on purified Pfs230c-dS/dS VLP derived from strain RK#114. Titration of VLP coating concentration. Primary antibodies were applied as 10µg/mL for both the mouse polyclonal and monoclonal 1B3). Error bars indicate standard deviation based on triplicates.

### S2.2 Anti-HCP Western blot

Anti-HCP Western blot was performed with the Pfs230c-dS/dS VLP preparation (Fig S 5, lane 1). The immunostaining of the membrane was performed as follows: Blocking with 1.5 % (w/v) powdered milk in PBS containing 0.05 % Tween 20 over-night at 4 °C. A polyclonal antiserum isolated from goats immunized with *H. polymorpha* HCP (Artes Biotechnology, Langenfeld, Germany/BioGenes, Berlin, Germany) was used as primary immunoreagent. The detection system was completed with a rabbit anti-goat IgG AP conjugate (BioRad, München, Germany) in combination with BCIP-NBT solution.

A subset of the bands detected in the Coomassie stained PAA gel (Fig S 5, lane 2) between the dS and the fusion protein was reactive with the polyclonal anti-HCP serum. Especially, the most prominent signals apart from the dS and the Pfs230c-dS in lane 2 could be identified as HCP in lane 1. Cross reactivity of the immunoreagents with the product-related proteins was not observed.

**Figure S 5.**
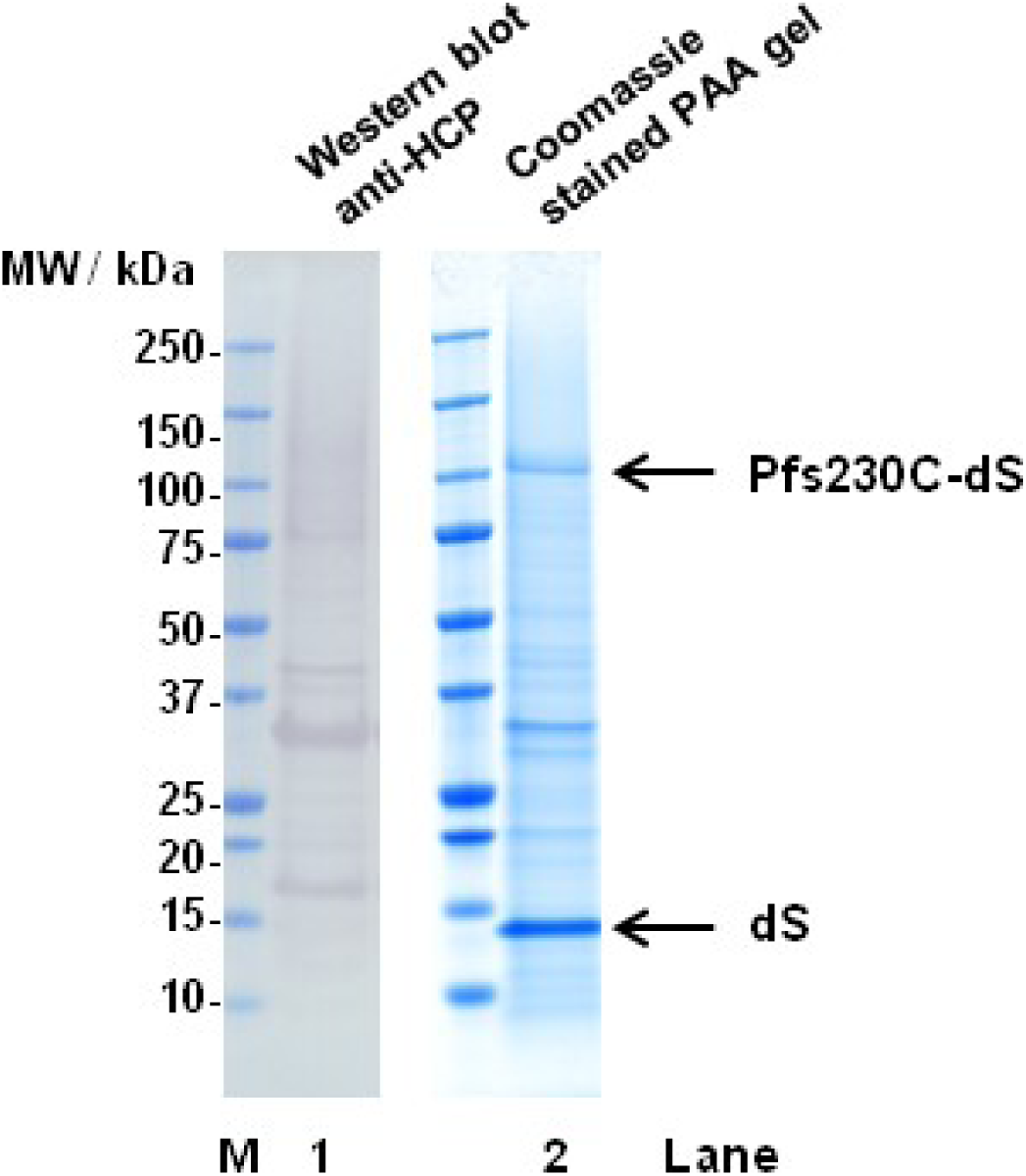
Anti-HCP Western blot analyses of purified Pfs230c-dS/dS VLP derived from strain RK#114. The purified Pfs230c-dS/dS VLP preparation was analyzed by Western blot probed with anti-HCP serum (lane 1, 10 µg protein loaded) or Coomassie stained PAA gel (lane 2, 12 µg protein loaded). M: molecular weight marker.

## Author contributions statement

Conceptualization, DW, JAC, JB, MP and DA; Methodology, DW, JAC, AB, EH, BK, CP, VJ, MW, MS, DA and MP; Investigation, DW, JAC, LR and AB; Administration, JB, MP, VJ, JR and DA; Writing – Original Draft, DW; Writing – Review and Editing, DW, JAC, MP, JB, MS, VJ and JM; Funding Acquisition, JB, MP and DA. Supervision, MP, JB, VJ, JM and GS.

## Funding

PATH Malaria Vaccine Initiative; National Health and Medical Research Council of Australia (Senior Research Fellowship and Program Grant to JB). Burnet Institute is supported by funding from the NHMRC Independent Research Institutes Infrastructure Support Scheme and a Victorian State Government Operational Infrastructure grant.

ARTES Biotechnology GmbH provided support in the form of salaries for authors DW, MS, MW, VJ and MP. MP is founder of ARTES Biotechnology GmbH and Managing Director and was involved in the study design, data analysis, decision to publish, and preparation of the manuscript as a supervisor as articulated in the ‘author contributions’ section. Most of the research presented in this work was performed in the company’s facilities.

Juliane Merz is employed by Evonik Technology & Infrastructure GmbH. Evonik Technology & Infrastructure GmbH provided support in the form of salary for author JM, but did not have any additional role in the study design, data collection and analysis, or preparation of the manuscript. Evonik Technology & Infrastructure GmbH expressly agreed on publishing the manuscript. The specific roles of JM are articulated in the ‘author contributions’ section.

## Acknowledgements

The authors gratefully acknowledge Sylvia Denter, Paul Gilson, Heribert Helgers, Renske Klassen, Dr. Andreas Kranz, Christine Langer, Dr. Joachim Leitsch, Thomas Rohr, and Elisabeth Wesbuer for technical and academic assistance. The following reagent was obtained through BEI Resources NIAID, NIH: Monoclonal Antibody 4B7 Anti-Plasmodium falciparum 25 kDa Gamete Surface Protein (Pfs25), MRA-28, contributed by David C. Kaslow. Monoclonal antibody 32F81 was kindly provided by PATH Malaria Vaccine Initiative. Polyclonal mouse antibody was kindly provided by Carole Long and Kazutoyo Miura, NIH.

## Competing interests

The authors VJ, MP, MS, DW and MW are associated with ARTES Biotechnology GmbH which owns the license for the VLP technology (patents cited as references [41] and [42]): Viral vectors expressing fusion of viral large envelope protein and protein of interest (No. WO2004092387A1). Recombinant proteins and virus-like particles comprising L and S polypeptides of avian hepadnaviridae and methods, nucleic acid constructs, vectors and host cells for producing same (No. WO2008025067A1).

Author JM is affiliated with Evonik Technology & Infrastructure GmbH.

There are no further patents, products in development or marketed products to declare. This does not alter our adherence to all the PLOS ONE policies on sharing data and materials.

## Data Availability

All relevant data are within the paper.

